# Microbial aerotrophy enables continuous primary production in diverse cave ecosystems

**DOI:** 10.1101/2024.05.30.596735

**Authors:** Sean K. Bay, Gaofeng Ni, Rachael Lappan, Pok Man Leung, Wei Wen Wong, Sophie Holland, Nadeesha Athukorala, Kalinka Sand Knudsen, Ziqi Fan, Melina Kerou, Surbhi Jain, Oliver Schmidt, Vera Eate, David A. Clarke, Thanavit Jirapanjawat, Alexander Tveit, Tim Featonby, Susan White, Nicholas White, Melodie A. McGeoch, Caitlin M. Singleton, Perran L.M. Cook, Steven L. Chown, Chris Greening

## Abstract

Most aerated cave ecosystems are assumed to be oligotrophic given they receive minimal inputs of light energy. Diverse microorganisms have nevertheless been detected within caves, though it remains unclear what strategies enable them to meet their energy and carbon needs. Here we determined the processes and mediators of primary production in aerated limestone and basalt caves through paired metagenomic and biogeochemical profiling. Based on 1458 metagenome-assembled genomes, over half of microbial cells in caves encode enzymes to use atmospheric trace gases as energy and carbon sources. The most abundant microbes in these systems are chemosynthetic primary producers, notably the novel gammaproteobacterial methanotrophic order *Ca*. Methylocavales and two uncultivated actinobacterial genera predicted to grow on atmospheric hydrogen, carbon dioxide, and carbon monoxide. *In situ* and *ex situ* biogeochemical and isotopic measurements consistently confirmed that these gases are rapidly consumed at rates sufficient to meet community-wide energy needs and drive continual primary production. Conventional chemolithoautotrophs, which use trace lithic compounds such as ammonium and sulfide, are also enriched and active alongside these trace gas scavengers. These results indicate that caves are unique in both their microbial composition and the biogeochemical processes that sustain them. Based on these findings, we propose caves are the first known ecosystems where atmospheric trace gases primarily sustain growth rather than survival and define this process as ‘aerotrophy’. Cave aerotrophy may be a hidden process supporting global biogeochemistry.

## Introduction

Found beneath one fifth of land surfaces, terrestrial caves provide unique habitats for life, given that they are dark, humid, thermally insulated, and relatively isolated systems^1–3^. The interiors of caves are typically oligotrophic habitats because, with specific exceptions (e.g. seasonally flooded, anthropogenically lit caves), they contain minimal photosynthetically-derived organic matter. Transport of dissolved organic carbon in groundwater may support microbial communities^4^, though its refractory nature means it is of limited importance^5^. Despite this energy limitation, the sediments and mineral surfaces of caves harbour abundant and diverse microbiota. Caves are enriched with the same nine dominant bacterial phyla as surface soils^6^, with many of these taxa thought to be aerobic organoheterotrophs^7– 10^. Some bacteria and archaea within caves are capable of harnessing chemical energy from reduced sulfur, nitrogen, and iron compounds present in drip water and mineral surfaces^11–14^. Given that these lithic compounds generally occur in trace amounts, chemosynthetic processes are thought to play a minor role in cave microbial ecosystems, except in globally rare, deep geothermally-heated caves usually isolated from the surface^15–20^.

A potential alternative source of energy and carbon in typical cave ecosystems is the atmosphere itself. We have recently discovered that atmospheric molecular hydrogen (H_2_) and carbon monoxide (CO) are critical energy sources supporting the biodiversity of soils and waters worldwide, and enable complex ecosystems to form in oligotrophic environments such as Antarctic soils^21–25^. Bacteria use high-affinity hydrogenases and CO dehydrogenases to liberate electrons from these gases for aerobic respiration and carbon fixation *via* the Calvin-Benson-Bassham (CBB) cycle^26^. Aerobic methanotrophs, which use atmospheric methane (CH_4_) as a dual energy and carbon source^27–29^, have also been identified in various cave systems and mediate CH_4_ oxidation rates comparable to those of surface soils^30–40^. Considering these findings, we sought to disentangle the relative roles of atmospheric, lithic, and solar energy sources in supporting primary production and energy conservation in cave ecosystems. To do so, we integrated genome-resolved metagenomic profiling, *in situ* and *ex situ* biogeochemical and isotopic measurements, and thermodynamic modelling of sediment and biofilm microbial communities collected along transects from four aerated limestone and basalt caves sampled within Australia.

## Results and Discussion

### Most cave microbes encode enzymes to harvest atmospheric energy sources

The samples from the four caves we sampled (Fig. 1a, Extended Data Table 1) spanned a broad range of organic carbon (0.08-28.4%), pH (3.6-8.7), and moisture levels (10.8-55.8%). Based on shotgun metagenomic profiling, microbial communities varied substantially between cave sediments and biofilms, between lithology types, and with cave depth (Fig. 1b & 1c; Extended Data Table 3a-g). Microbial abundance (av. 6.7 × 10^9^ rRNA gene copies per gram of dry sediment) and richness (av. 582 Nearest Taxonomic Units per sample; Chao1 (based on metagenomic 16S rRNA gene) decreased by 3.3-fold and 1.5-fold, respectively, between the cave entrance and interior sediments (Fig. 1; Extended Data Table 2 & 3b). In line with most other sampled caves^9^, Actinobacteriota, Proteobacteria, Acidobacteriota, Chloroflexota, and Gemmatimonadota were the most common phyla, along with Thermoproteota (predominantly Nitrosophaerales) (Fig. 1b & 1c; Extended Data Table 3d). Metagenomic assembly and binning yielded 1458 dereplicated high- and medium-quality metagenome-assembled genomes (MAGs) spanning 36 different phyla.

**Figure 1.**
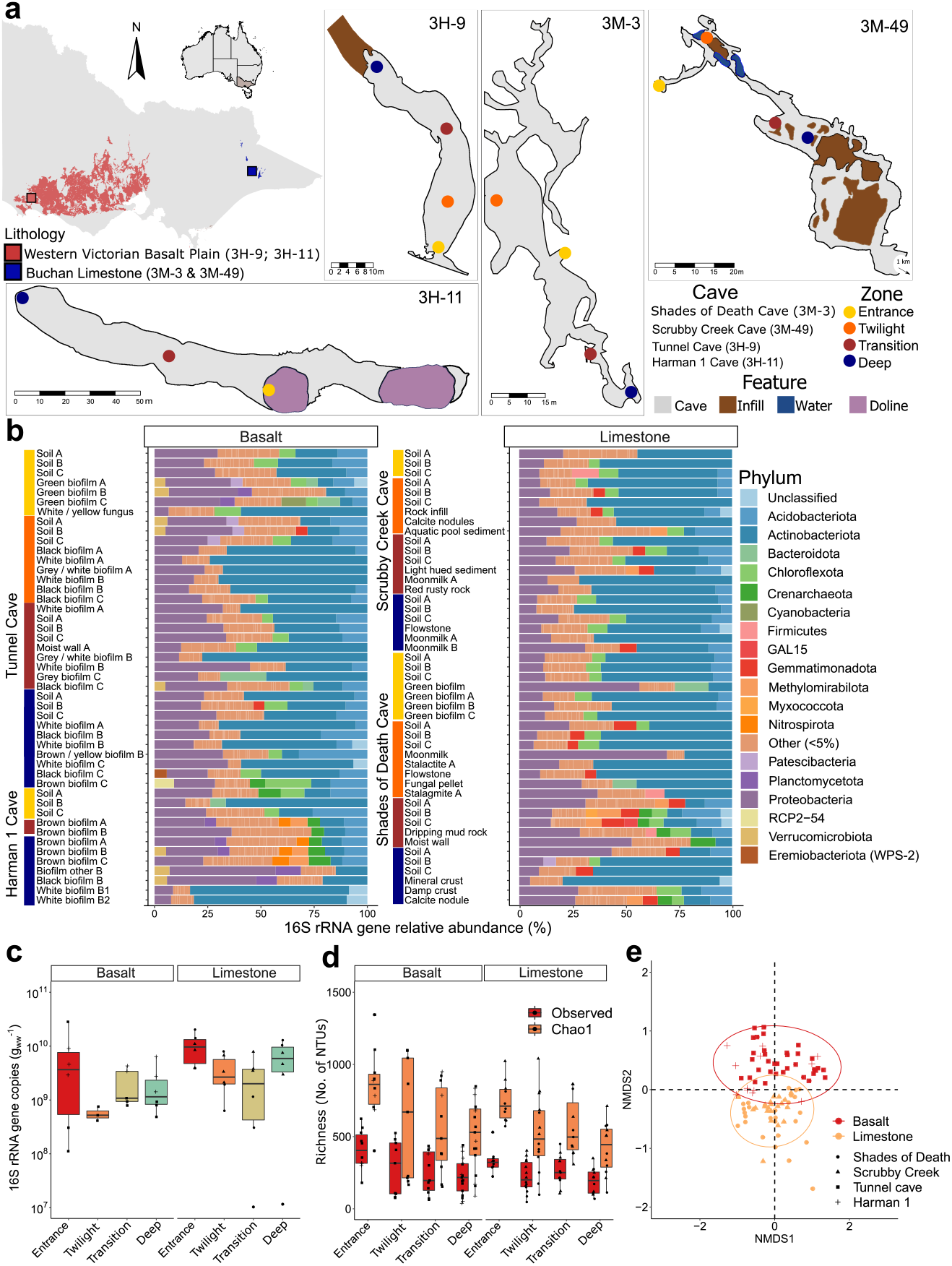
Cave microbial community composition and structure. **a**, Map showing geographic and lithological setting of the caves studied. Detailed cave morphology, major environment features, study sites, scale, and sites are shown. **b**, Phylum-level community composition at the sample level, vertical-coloured bars show the environmental diversity captured according to sample types such as biofilms and soils. **c**, Boxplot of 16S rRNA gene copy number for bulk sediments and peanut butter biofilms according to site and host lithology. **d**, Boxplot of observed and estimated richness according to site and host lithology. **e**, Non-metric multidimensional scaling using Bray-Curtis similarity comparing differences in community structure according to lithology.

Inferring the energy and carbon acquisition strategies of the cave microbes by searching for 52 conserved marker genes in the MAGs and short reads (Fig 2a; Extended Data Table 4a-c) suggested that most cave bacteria mediate aerobic respiration using both organic compounds and trace gases as substrates (Fig. 2a-b). Numerous MAGs (44%, normalised to genome completeness), accounting for 54% of mapped metagenomic reads (Extended Data Table 4c), encoded enzymes to consume one or more atmospheric trace gases, namely form I CO dehydrogenases for CO oxidation (25.4% genomes / 73% community based on short reads; Fig. 2a-b; Extended Data Fig. 1), group 1 and 2 [NiFe]-hydrogenases for H_2_ oxidation (25.8% / 43%; Fig. 2a-b; Extended Data Fig. 2), and particulate methane monooxygenases for CH_4_ oxidation (2.9% / 5.1%; Fig. 2a-b; Extended Data Fig. 3a-c). These findings suggest that most microbial cells in caves can oxidise trace gases. Many cave microbes can also use lithic energy sources such as sulfide (11.3% / 19%; Fig. 2a-b), thiosulfate (6.6% / 7.3%, Fig. 2a-b), ammonia (4.1% / 3.6%; Extended Data Fig. 4), nitrite (1.2% / 3.7%; Extended Data Fig. 5), and ferrous iron (2.7% / 2.3%; Fig. 2a-b). Photosynthesis genes were abundant in sediments and biofilms at the entrance of each cave, but declined by an average of 75-fold in the cave interior. Conversely, we observed enrichment in cave interiors compared to the entrance of the genes enabling the oxidation of CH_4_ (7-fold), ammonium (1.9-fold), nitrite (2.3-fold), sulfide (1.3-fold), and to a lesser extent, H_2_ (1.3-fold), and CO (1.3-fold) oxidation. Concordant patterns were observed for carbon fixation genes, with the photosynthetic cyanobacterial type IB RuBisCO decreasing 38-fold, and the chemosynthetic, predominantly actinobacterial type IE RuBisCO increasing 2-fold in cave interiors compared to entrances (Extended Data Table 4b & c; Fig. 2a & b; Extended Data Fig. 6). Some carbon fixation also likely occurs through 4-hydroxybutyrate cycle (2.7% / 2.4%), reductive tricarboxylic acid cycle (1.2% / 2.2%), and 3-hydroxypropionate cycle (0.6% / 0.7%). Altogether, these findings indicate a shift from photosynthetic to chemosynthetic primary production in cave ecosystems, driven both by atmospheric and lithic substrates.

**Figure 2.**
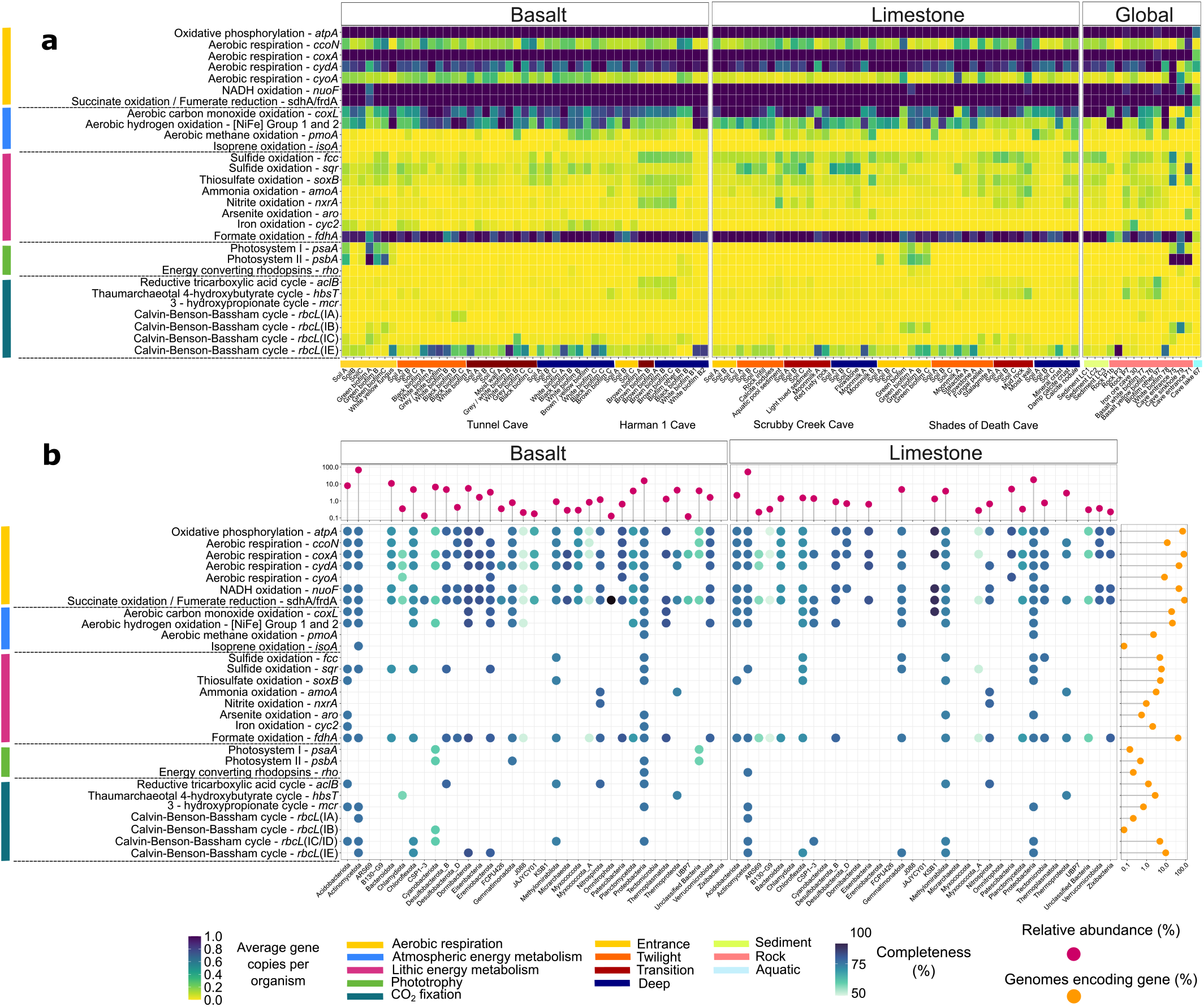
Metabolic potential of energy and carbon acquisition. **a**, Heatmap showing the metabolic potential of the community as average gene copies per organisms for conserved marker genes of major energy and carbon acquisition pathways across limestone and volcanic caves in Australia in comparison to global samples. **b**, Metabolic potential at the genome resolved level (MAGs). Each dot in the blue shading represents the presence of encoded metabolic functions and the shading represents average genome completeness at the phylum level. Orange lollypop charts show the percentage of genes encoded across all MAGs and purple lollypop charts show the relative abundance maximum for each phylum.

**Figure 3.**
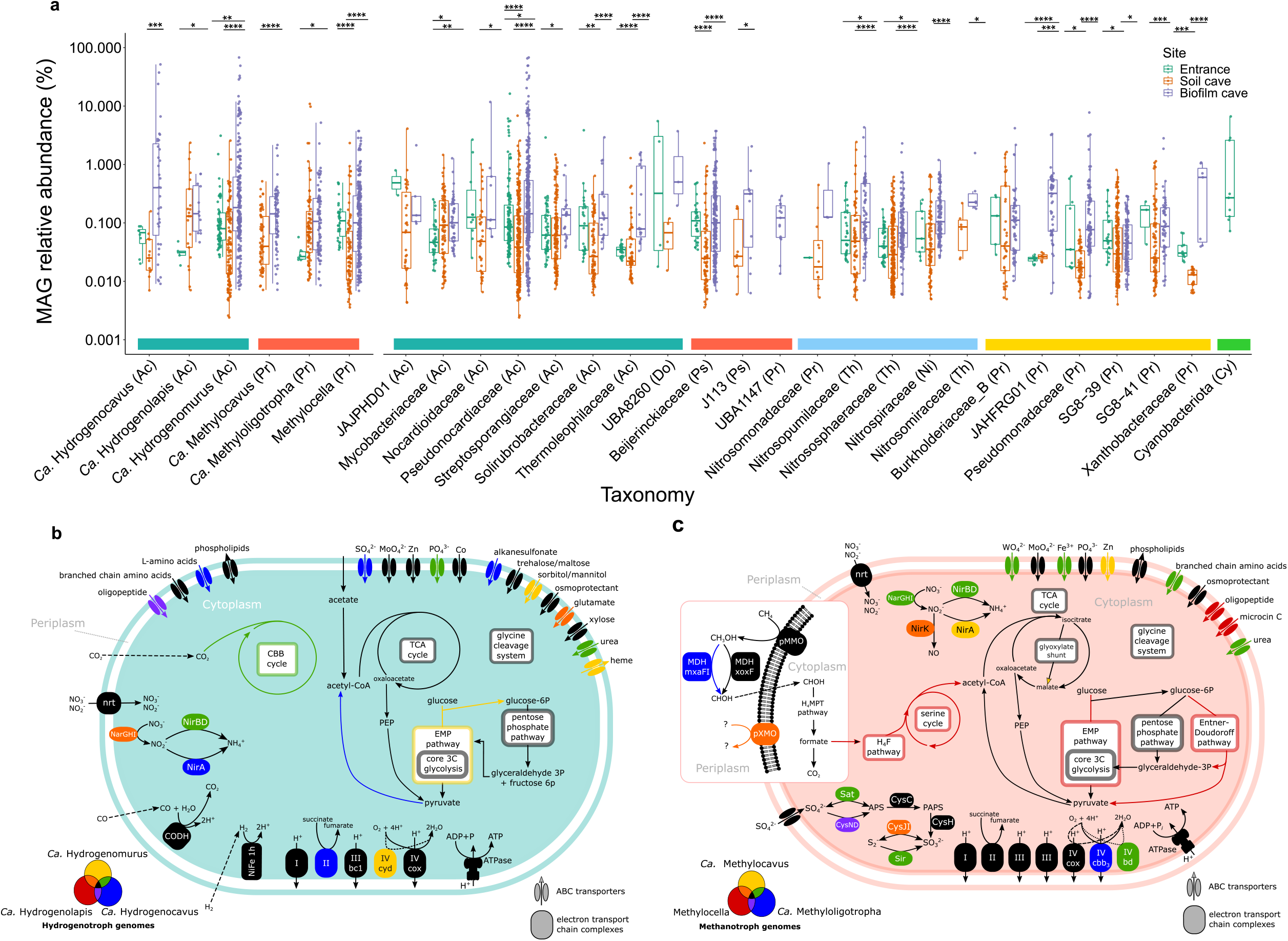
Abundance and capabilities of the most abundant functional groups in caves. **a**, Differential abundance of key taxa, at family level, between cave entrance, interior sediment, and interior biofilm samples. Box plots show the relative abundance, based on read mapping, of MAGs of hydrogenotrophs (teal), methanotrophs (red), nitrifiers (blue), sulfur oxidisers (yellow), and phototrophs (green). Pairs are denoted with asterisks showing significant enrichment. Taxonomic classification is shown at genus level for taxa used for metabolic mapping. Except for Cyanobacteriota, taxa are shown at family level and at its preceding rank if unclassified, with brackets showing phylum level affiliation (Ac – Actinobacteriota; Pr - Proteobacteria; Do – Dormibacteriota; Th – Thermoproteota; Ni – Nitrospirota). **b**, Metabolic reconstruction of the three dominant hydrogenotrophic taxa, the candidate genera *Hydrogenomurus, Hydrogenocavus*, and *Hydrogenolapis*. All encode genes consistent with trace atmospheric gas oxidation, including a group 1h [NiFe] hydrogenase and CO dehydrogenase. **c**, Metabolic reconstruction of the three dominant methanotrophic MAGs, *Methylocella* (USCα), Methyloligotrophales (USCγ), and Methylocavales. All encode particulate methane monooxygenase and a methanol dehydrogenase. Carbon fixation in *Methylocella* MAGs can occur via the tetrahydrofolate pathway and serine cycle, but remains unresolved in the Gammaproteobacterial MAGs.

**Figure 4.**
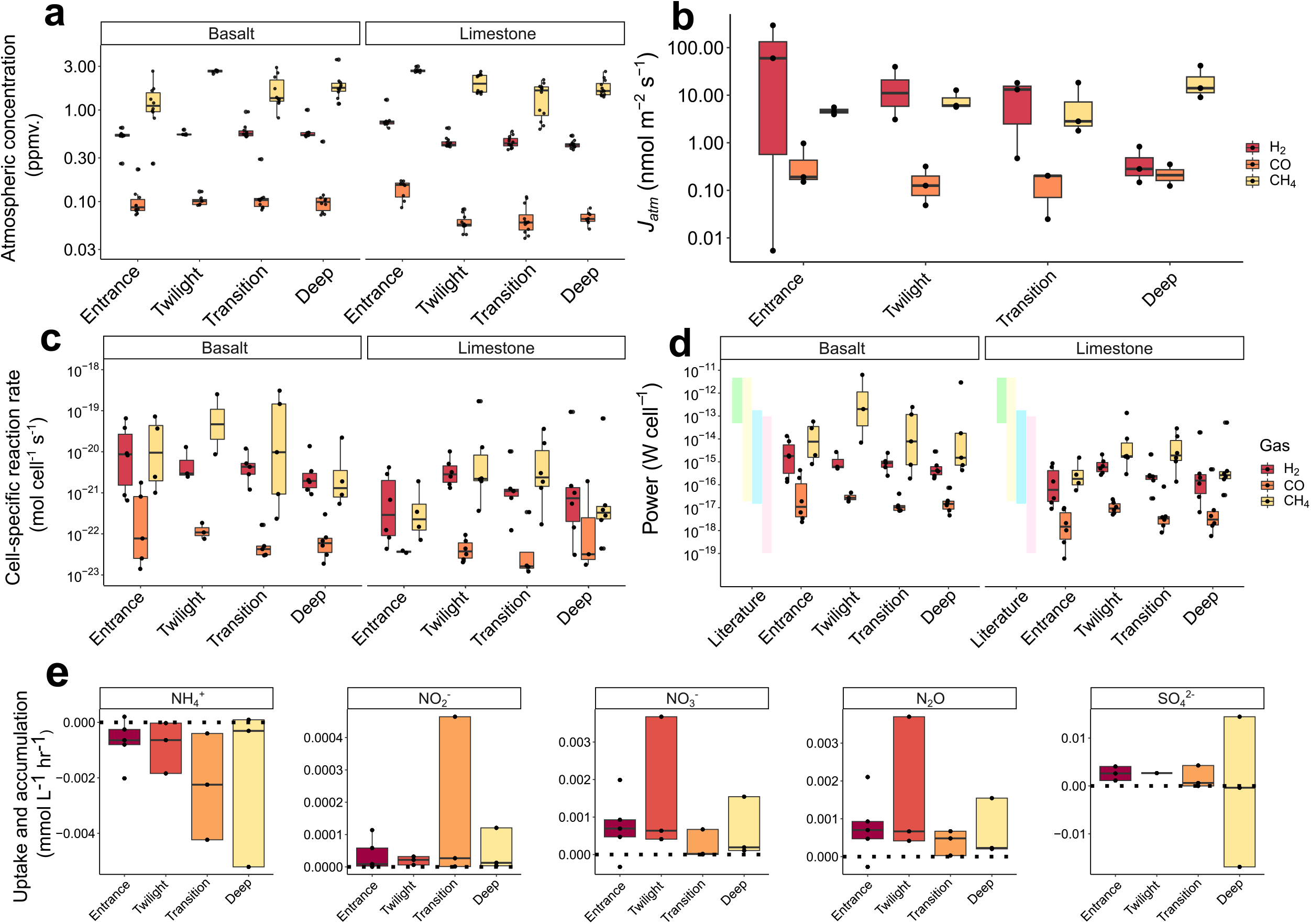
*In situ, ex situ* and energy yield measurements for trace gases H_2_, CO and CH_4_. **a**, *In situ* atmospheric concentrations (ppmv.) for all four caves, faceted by each of the three gasses and site. **b**, *In situ* sediment–atmosphere gas fluxes (*J*_*atm*_ negative values indicate net gas consumption). **c**, Bulk sediment oxidation rates over time faceted by each gas and site. **d**, Amount of power per cell derived from the oxidation of each trace gas, coloured bars depict the range of literature values of maintenance energy requirements or endogenous metabolic rates of different pure cultures (green^61^, yellow^62^, lilac^63^) and hydrogen oxidisers in deep marine sediments (pink^62,64^). **e**, Rates of nitrogen and sulfur compound metabolism, with positive values indicating accumulation and negative values showing uptake. Nitrification is expected to result in ammonium (NH_4_ ^+^) consumption and nitrite (NO_2_^-^), nitrate (NO_3_^-^), and nitrous oxide (N_2_O) production, whereas sulfide oxidation is expected to cause sulfate (SO_4_^2-^) production. All boxplots show min., max., median, and IQR.

**Figure 5.**
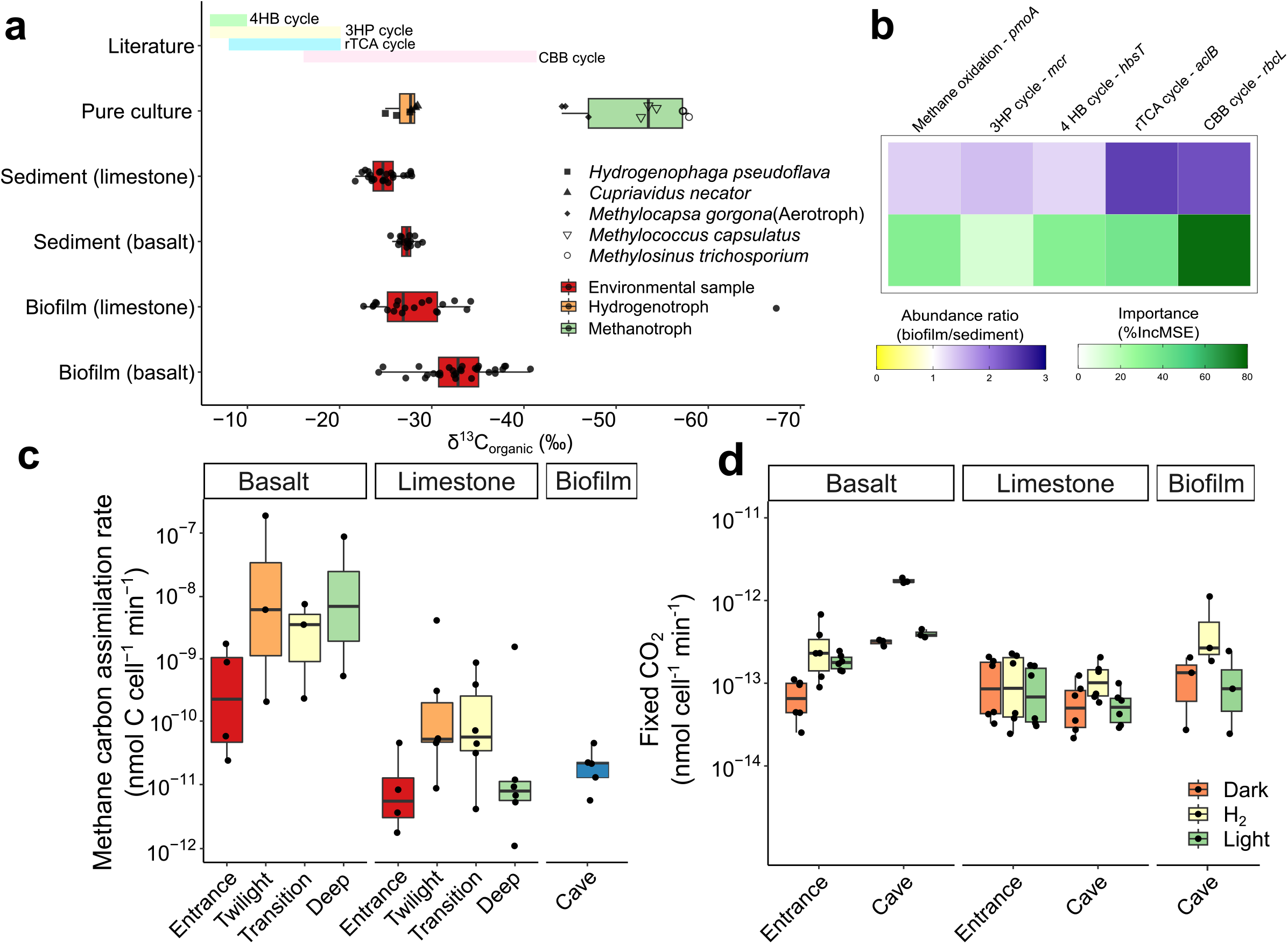
Major carbon acquisition processes and activities in caves. **a**, Boxplot showing depletion of biomass ^13^C stable isotope (δ^13^C_organic_) across cave sediments, biofilms, and selected autotrophically-grown hydrogenotroph and methanotroph pure cultures. Coloured bars depict the range of literature values of δ^13^C_organic_ of biomass produced from 4-hydroxybutyrate cycle (4HB cycle; green), 3-hydroxypropionate cycle (3HP cycle; yellow), reductive tricarboxylic acid cycle (rTCA; blue), and Calvin-Benson-Bassham cycle (CBB cycle; pink). δ^13^C_organic_ for pure cultures and literature values were adjusted based on the use of atmospheric CO_2_ (δ^13^C: −8.5‰) and CH_4_ (δ^13^C: −47.2‰) as sole carbon sources. **b**, Heatmap showing the abundance ratio of key carbon assimilation marker genes (*pmoA, mcr, hbsT, aclB, rbcL*) in biofilm against sediment communities (top) and random forest analysis (% Mean Squared Error) of these genes as predictors for cave δ^13^C_organic_ values (bottom). **c**, Boxplot showing methane carbon assimilation rate as a function of cellular oxidation rates normalised by median methane carbon assimilation fraction commonly observed across soil ecosystems. **d**, Cellular ^14^C-CO_2_ fixation rates faceted by lithology, comparing cave entrance with interior across three conditions.

To ensure these insights were representative of caves worldwide, we further analysed twelve previously published metagenomes representative of diverse global cave ecosystems (Fig. 2a, Extended Data Table 4a). Oxidation of ambient trace gases and, to a lesser extent, lithic inorganic compounds are widespread strategies in sediments and rocks of cave interiors. For example, in rock metagenomes from Monte Cristo Cave (Brazil) and in white microbial mats in Kipuka Kanohina Cave (Hawaii), almost all microbes encode high-affinity hydrogenases^41^. The three exceptions are photosynthetic biofilms collected from an illuminated entrance and sinkhole, as well as a cave lake likely to receive considerable organic inputs (Extended Data Table 4a).

### Novel microbes drive cave energy acquisition and primary production

We used genome-resolved metagenomics and phylogenetic analyses to resolve which microbes mediate these processes (Extended Data Table 4c; Extended Data Fig. 1 to 6). Most hydrogenases, CO dehydrogenases, and RuBisCOs were co-encoded by the most abundant Actinobacteriota lineages residing in the caves (primarily classes Actinomycetia, Thermoleophilia, Acidimicrobiia, and *Ca*. Aridivitia)^23^ (Fig. 3a), suggesting that they are the dominant primary producers in these ecosystems. Multiple phyla nevertheless encoded each of these enzymes (12 hydrogenase-, 11 CO dehydrogenase-, and 11 RuBisCO-encoding phyla), highlighting that trace gas oxidation and chemosynthesis are ubiquitous traits (Extended Data Table 4). These enzymes were also encoded by various uncultivated lineages, for example with CO dehydrogenases being encoded by high-quality genomes from the candidate bacterial phyla CSP1-3 and KSB1, as well as two enigmatic orders (RBG-16-68-12, UBA184) of Thermoplasmata archaea inhabiting diverse cave samples (Extended Data Fig. 1). Corroborated by the short-read analysis (Fig. 2a, Extended Data Table 4b), almost all of these hydrogenases are high-affinity clades (groups 1h, 1l, and 2a [NiFe]-hydrogenases^23,42,43^) (Fig. 2a), highlighting adaptation to atmospheric concentrations rather than higher concentrations of these gases. Of the 30 most abundant microbes based on genome read mapping (Extended Data Fig. 3a-b, Extended Data Table 4c), 21 were capable of trace gas oxidation including four methanotrophs (all affiliated with the USCγ / JACCXJ01 clade), whereas none mediated photosynthesis, nitrification, sulfide oxidation, or iron oxidation. The top ten most abundant microbes (comprising 7.6% of all reads) were all from uncultivated genera from Pseudonocardiaceae and Egibacteraceae (both within class Actinomycetia), each of which co-encoded RuBisCO with either CO dehydrogenase and/or uptake hydrogenases. This indicates that caves select highly productive actinobacterial primary producers that grow on atmospheric energy and carbon sources. These Actinobacteriota are the most abundant lineages in the cave biofilms, whereas the methanotrophs are the single most abundant species in the cave sediments.

We comprehensively analysed the energy and carbon acquisition pathways of the three most abundant predicted hydrogenotrophs in the caves (Fig. 3b), namely the candidate genera herein named *Hydrogenomurus, Hydrogenocavus*, and *Hydrogenolapis* (all etymological information in **Supplementary Note 2**; formerly Pseudonocardiaceae GCA-003244245, Egibacteraceae JACCXR01, and Actinomycetia JACCUZ01). These lineages were selectively enriched in distinct niches, with *Hydrogenomurus* prevalent across basalt caves and constituting over half of multiple biofilm communities (up to 73%), *Hydrogenocavus* dominant in limestone biofilms and moonmilks (up to 54%), and *Hydrogenolapis* abundant in limestone biofilms and sediments (Fig. 3a; Extended Data Table 4c). All three taxa encoded high-affinity group 1h [NiFe]-hydrogenases and CO dehydrogenases, consistent with use of trace gases as an energy source. In addition, these MAGs encoded a complete TCA cycle and a near complete suite of aerobic respiratory complexes (I-V), indicating they can conserve energy through both lithotrophic and organotrophic aerobic respiration. Both *Hydrogenomurus* and *Hydrogenocavus* also encoded type IE RuBisCO and a complete CBB cycle, indicating they are facultative chemolithoautotrophs; the lack of enzymes for oxidation of lithic compounds strongly suggests reductants necessary for carbon fixation are provided by H_2_ and CO. Their autotrophic capacity likely underlines their dominance along oligotrophic cave mineral surfaces. Conversely, *Hydrogenolapis* appears to be solely reliant on organic carbon sources, potentially including peptides and mono- and disaccharides, based on the presence of various ABC transporters. Energy provided by atmospheric trace gases likely enables these microbes to allocate more organic carbon for anabolism^44^. All taxa encoded the pentose phosphate and Embden–Meyerhof–Parnas pathways for organic carbon catabolism, although only *Hydrogenomurus* MAGs encoded all genes necessary for complete glycolysis from glucose.

Methanotrophs were the most enriched metabolic specialists in the cave interior, based on both genomic read mapping (Fig. 3a; Extended Data Table 4b) and marker gene profiles (Fig. 2a & b). Therefore, we performed an in-depth analysis to resolve the evolutionary history and functional capabilities of putative cave methanotrophs encoding particulate methane monooxygenases (pMMO), as elaborated in **Supplementary Note 1**. A genome tree revealed that these bacteria span the alphaproteobacterial genus *Methylocella* (encompassing *Methylocapsa* within the GTDB framework), the gammaproteobacterial order Methylococcales, and two candidate gammaproteobacterial orders herein named Methyloligotrophales and Methylocavales (formerly JACCXJ01 and CAJXQU01; etymology in **Supplementary Note 2**) (Extended Data Fig. 3a). These methanotrophs were progressively enriched from entrance and cave sediments to biofilms, with a single Methyloligotrophales MAG encompassing 10.8% of microbes in a complex limestone sediment (Fig. 3a; Extended Data Table 4c), suggesting CH_4_ is a primary growth substrate of cave communities. While *Methylocella* and Methyloligotrophales encompass the USCα and USCγ lineages of atmospheric methanotrophs^27,28,45,46^, Methylocavales is not known to be methanotrophic and is represented by just one previously reported genome that lacks *pmo* genes. Consistent with being novel methanotrophs, the cave-exclusive Methylocavales bacteria each encoded complete *pmoCAB* operons and their PmoA protein formed a novel clade sister with Methyloligotrophales (Extended Data Fig. 3b). They also encode a complete set of genes to oxidise methanol (lanthanide-dependent methanol dehydrogenases), formaldehyde (tetrahydromethanopterin pathway), and formate (formate dehydrogenase) to carbon dioxide for energy conservation (Extended Data Fig. 3c). However, in common with other atmospheric gammaproteobacterial methanotrophs, including the Methyloligotrophales MAGs analysed, the carbon assimilation pathways used remain incompletely resolved (**Supplementary Note 1**).

Although the capacity for oxidation of lithic substrates was less widespread, numerous chemolithoautotrophs were nevertheless highly enriched in cave sediments and biofilms. Most notable are nitrifiers, including ammonia-oxidising archaea and bacteria, as well as nitrite-oxidising and comammox Nitrospirales (Fig. 2a & b; Extended Data Table 4b-c). Members of three archaeal families, Nitrososphaeraceae, Nitrosopumilaceae (e.g. acidophilic *Nitrosotalea* dominant in basalt caves), and novel clade Nitrosomiraceae (e.g. *Ca*. Nitrosomirus^47^ abundant in limestone caves), vastly outnumber ammonia-oxidizing bacteria (*Nitrosospira*) (Fig. 3a); most encode high-affinity Amt1 ammonia transporters, carbon sequestering ABC-type bicarbonate transporters, and carbonic anhydrase consistent with their oligotrophic lifestyle^48^. Remarkably, urease, cyanase, and glycine cleavage system genes were also present in most MAGs, suggesting these archaea also sequester ammonia from organic substrates such as urea, cyanate, and glycine. Comammox Nitrospirales (genus Palsa-1315), which include lineages known for their high affinity for ammonia^49^, were also widely distributed (Extended Data Table 4b-c). These cave nitrifiers use distinct pathways to fix CO_2_, spanning 4-hydroxybutyrate cycle (Nitrososphaerales), reductive tricarboxylic acid cycle (Nitrospirales), and CBB cycle (*Nitrosospira*). Altogether, these findings indicate that caves select for oligotrophic chemolithoautotrophs, and that nitrifiers are likely important primary producers given their autotrophic lifestyle and enrichment. Apart from nitrification, some nine phyla were capable of sulfide oxidation, including numerous Proteobacteria and Actinobacteriota MAGs. We also reconstructed genomes of iron-oxidising Acidobacteriota and Proteobacteria (Fig. 2a & b).

### Trace gas oxidation occurs at high rates alongside lithic substrate oxidation

To substantiate these findings, we performed *in situ* and *ex situ* profiling of the processes of trace gas oxidation and lithic substrate oxidation in each cave. *In situ* measurements of ambient average concentrations of CH_4_, H_2_ and CO at the cave entrance were 1.89, 0.63 and 0.11 ppmv, respectively (Fig. 4a; Extended Data Fig. 7), which reflect similar global average concentration of these gases in the lower troposphere^50–52^. Ambient air concentrations of CH_4_ and H_2_ gases in limestone caves decreased 1.6-fold and 1,8-fold respectively from the cave entrance to the interior, suggesting microbial consumption (Fig. 4a; Extended Data Fig. 7). *In situ* CH_4_ fluxes greatly increased from −4.7 nmol m^−2^ s^−1^ at the entrance to an average of −37 nmol m^−2^ s^−1^ inside, confirming vast methanotroph activity within caves (Fig. 4b; Extended Data Fig. 7, Extended Data Table 5a). In contrast, H_2_ fluxes were highest at the entrance and modestly declined inside the cave, averaging around −25 nmol m^−2^ s^−1^ (Fig. 4b; Extended Data Fig.7, Extended Data Table 5a).

Given the limitations in conducting flux measurements only in areas with sufficient sediment depth for flux chambers, we employed microcosm incubations with bulk sediments and biofilms extracted from cave walls to validate these observations. The microbes within cave sediments and biofilms rapidly consumed all three gases to below atmospheric concentrations (Fig. 4c; Extended Data Fig. 8, Extended Data Table 5b). Oxidation rates were highest for H_2_ on average, followed by CH_4_ and CO. Notably, *ex situ* CH_4_ oxidation rates closely matched *in situ* patterns, especially in limestone caves, with a remarkable 7-fold increase from entrance to deep zones (Fig. 4c; Extended Data Fig. 8, Extended Data Table 5b), in line with metagenomic observations of increased methanotrophic abundance (Fig. 2a). We also tested whether lithic substrates, namely ammonium and sulfide, were also used as energy sources given the metagenomic observations (Fig. 2a). The cave sediments contained varying concentrations of ammonium (1.35 – 40.2 mg/kg), sulfur (7.3 – 2468 mg/kg), and iron (6.96 – 2003 mg/kg) as potential chemical energy sources for chemolithoautotrophs (Extended Data Table 1). Ammonium was oxidised at variable rates across the samples and increased from the entrance to the cave interior (Fig. 4e; Extended Data Fig. 9; Extended Data Table 6). These incubations also revealed the accumulation of nitrite and nitrate, consistent with stepwise nitrification processes occurring within the cave environments. Sulfide oxidation was also evident from the accumulation of the end-product sulfate (Fig. 4e; Extended Data Fig. 9; Extended Data Table 6).

### Trace gas oxidation drives community-wide carbon and energy provision

Thermodynamic calculations based on bulk oxidation rates of H_2_, CO, and CH_4_ and cell estimates from metagenomics-adjusted qPCR quantification of 16S rRNA genes, revealed that trace gas oxidation rates yielded an average power output of 1.5 × 10^−15^, 3.7 × 10^−17^, and 2.6 × 10^−13^ W per H_2_, CO, and CH_4_-oxidising cell, respectively. Power per cell outputs were similar between lithology types and cave depths for H_2_ and CO, but for CH_4_ were significantly higher in basalt compared to limestone and between surface and subsurface (Fig. 4d; Extended Data Table 5b). For H_2_ and CO, these calculations are within the average range of maintenance energy reported for various isolates of cultured, typically copiotrophic aerobic organoheterotrophs (10^−12^ – 10^−17^ W cell^-1^)^53–55^ and exceed the theoretical maintenance at the limits of life (10^−17^ – 10^−19^ W cell^-1^)^56,57^. These rates are averaged for all cells and samples, although some microbes likely grow by rapidly co-consuming these gases, notably the highly abundant *Hydrogenocavus* and *Hydrogenomurus*. For methanotrophs, these rates greatly exceed the realm to support growth of recently cultivated atmospheric methanotrophs (1.9 × 10^−15^ W – 6.1 × 10^−16^ cell^-1^)^45,58^. Altogether, these models indicate that the rates of atmospheric trace gas consumption are sufficient to sustain the growth of the methanotrophs and the survival of the hydrogenotrophs in these caves, with some bacteria potentially also mediating chemolithoautotrophic growth by using atmospheric H_2_ and/or CO to fix CO_2_.

To probe the major pathways contributing to the organic matter in caves, we quantified the fractionation signature of biomass ^13^C/^12^C in cave sediments and biofilms (Fig. 5a). Two autotrophically-grown hydrogenotrophs and three methanotrophs (including atmospheric CH_4_ oxidiser *Methylocapsa gorgona*) were also analysed for their carbon fractionation as a comparison. Cave sample organic fractions display a depletion of ^13^C (δ^13^C_organic_) ranging from −21.7 to −67.4 ‰, consistent with patterns of hydrogenotrophs and biomass derived from CBB cycle but distinct from other carbon fixation pathways^59,60^ (Fig. 5a). δ^13^C_organic_ was progressively more negative from sediments in limestone caves to biofilms in basalt caves, in line with the increasing trends of *rbcL* and *pmoA* gene abundance (Fig. 2a, Fig. 5b). Random forest analysis reveals *rbcL* abundance is the best carbon assimilation gene predictor for δ^13^C_organic_ (Fig. 5b), and also the most negatively correlated with δ^13^C_organic_ among all metabolic marker genes (Spearman’s *rho* = −0.56, *p* = 2.0 × 10^−6^). This analysis suggests organic carbon in caves is predominantly derived from the CBB cycle. CH_4_ assimilation, which yields biomass strongly depleted in ^13^C, may also contribute to the highly negative δ^13^C_organic_ in some cave samples, such as a biofilm sample with a δ^13^C_organic_ of −67.4‰ (Fig. 5a). This was supported by methane carbon assimilation as a function of cell specific methane oxidation rates (Fig. 5c).

Finally, we traced radioisotope incorporation of ^14^C-CO_2_ into biomass to ascertain the relative contributions of dark, hydrogenotrophic, and photosynthetic CO_2_ fixation pathways, including as a source of the δ^13^C_organic_ signatures. Whereas photosynthesis was strongly stimulated in the entrance of basalt caves, it was negligible otherwise. Hydrogenotrophic CO_2_ fixation was observed in cave interiors, with two-fold and five-fold more carbon fixed in limestone and basalt caves respectively compared to dark conditions (Fig. 5d; Extended Data Table a). Notably, biofilm and basalt sediment microbes mediated particularly high hydrogenotrophic CO_2_ fixation activities (6.7 × 10^−12^ nmol cell^-1^ min^-1^) compared to average rates by limestone sediment microbes (6.9 × 10^−13^ nmol cell^-1^ min^-1^). This supports the metagenomic inferences that H_2_ is the predominant driver of CO_2_ fixation and likely the observed δ13C_organic_ signatures in caves. Methanotrophs also contribute significantly to carbon acquisition, as they assimilate 1.8 x 10^−7^ to 2.0 x 10^−12^ nmol C cell^-1^ min^-1^, which is further supported by their high activities based on the *in situ* flux analysis (Fig. 5b; Extended Data Table 7b). These experiments demonstrate that microbial energy and carbon acquisition from atmospheric substrates occur at significant rates with chemosynthetic primary productivity being continuously sustained across a range of cave surfaces given the relatively stable environmental settings. Aboveground, this type of chemosynthetic primary productivity typically exhibits greater flux variation due to environmental conditions such as aridity.

## Conclusions

Here we provide strong metagenomic and biogeochemical evidence that diverse caves are atmospherically-powered ecosystems. Primary production appears to be driven by highly abundant and active methanotrophs, as well as novel lineages of actinobacterial lithoautotrophs, that continuously use the gases methane, hydrogen, carbon dioxide, and carbon monoxide present in cave atmospheres. Cave ecosystems differ from polar desert soils, the other major type of ecosystem shown to be primarily atmospherically-powered^21–23^, in that trace gases appear to drive substantial continual growth rather than long-term survival in these nutrient-deprived environments. This is reflected by the abundant primary producers in cave sediments and biofilms, the rapid fluxes and activities of trace gas oxidisers, and the theoretical considerations based on thermodynamic and biogeochemical modelling. On this basis, we propose defining the term ‘aerotrophy’ as “the process of growth through the use of atmospheric trace gases as energy and carbon sources” and redefine caves as ecosystems often driven by ‘aerotrophic microorganisms’. Nevertheless, there is much spatial variation in the mediators and rates of this process across cave ecosystems. Other energetic processes co-occur, including chemolithoautotrophic nitrification and sulfide oxidation, with these processes likely becoming dominant in environments where these substrates are more abundant. Aerotrophy might not be as dominant in the numerous caves that have more extensive solar or organic carbon inputs, are disconnected from the atmosphere, or are otherwise largely anoxic. Nonetheless, the occurrence of caves beneath 20% of ice-free terrestrial areas suggests that aerotrophy likely supports large and diverse ecosystems worldwide. Cave aerotrophy may thus be a hidden process influencing global biogeochemical cycling of hydrogen, methane, and carbon.

## Footnotes

## Acknowledgements

This research was conducted as part of the Australian Research Council (ARC) SRIEAS grant Securing Antarctica’s Environmental Future (SR200100005; awarded to S.L.C., C.G., M.A.M.) and a Monash University Faculty of Science Strategic Uplift Seed Grant (Awarded to S.K.B., P.C., W.W.W.). S.K.B. and R.L. are supported by an ARC Discovery Early Career Research award (DE230101346; DE230100542). C.G. is supported by an NHMRC EL2 Fellowship (APP1178715). C.M.S. is supported by a Novo Nordisk Foundation Postdoctoral Fellowship grant (NNF20OC0065005). K.S.K. is supported by the DarkScience project (Villum Foundation). G.N. and P.M.L. acknowledge the Early Career Postdoctoral Fellowship (ECPF23-8566329039 & ECPF23-1113137961) awarded by Faculty of Medicine, Nursing and Health Science (FMNHS) at Monash University. A.T.T. and O.S. were supported by the Research Council of Norway (projects Living on Air, 315129 and Harvesting Energy from Air, 347122). Z.F. is supported by Monash University-China Scholarship Council Joint Scholarship (CSC202308240008). This study used the MASSIVE M3 and MonARCH supercomputing infrastructure. We are grateful of Tess Hutchinson and Michaela Wawryk for technical support.

## Author contribution

S.K.B., C.G., and S.L.C conceived this study. S.K.B. planned and led field work, designed and led experiments and analysed data. Different authors were responsible for performing fieldwork (S.K.B., R.L., T.F., P.M.L., S.W., N.W.), field logistics (S.K.B., S.W., T.F., N.W.), DNA extraction and qPCR (S.K.B., R.L., T.J.), gas chromatography (S.K.B.), nitrification and sulfide oxidation assays (W.W.W., V.E., P.M.L.C.), pure culture preparation (N.A., S.J., O.S., A.T.), carbon stable isotope measurement (W.W.W., P.M.L., Z.F., V.E., P.M.L.C.), ^14^C radioisotope tracing (S.K.B.), spatial analysis and mapping (D.A.C., T.F., S.W., M.A.M.), metagenomic community and metabolic analysis (S.K.B., G.N., C.G. P.M.L.), MAG construction and annotation (G.N.), and metabolic reconstruction (S.H., K.S.K., N.K., M.K, C.M.S., S.K.B., G.N., C.G., P.M.L.). S.L.C. and C.G. provided most resourcing, supervision, and funding. S.K.B., C.G., S.L.C., and P.M.L. wrote the paper with input from all authors.

## Data Availability Statement

All previously sequenced metagenomes analysed in this study are available at NCBI BioProject with the accession numbers listed in Supplementary Data Table 4a. All metagenomes sequenced for this project are deposited at the NCBI Sequence Read Archive PRJNA1048116. All metagenome assembled genomes are available at https://figshare.com/s/80196efcf5886c767713 and will be published to GenBank prior to publication.

## Ethics declarations

The authors declare no competing financial interests.

