## Supplementary Note for "Microbial aerotrophy enables continuous primary production in diverse cave ecosystems"

**This file includes:**

- Methods
- Extended Data Figures 1-9
- Extended Data Tables 1-7
- Supplementary note 1 and 2

### **Methods**

#### **Site description and sampling**

Samples were collected from four caves in Victoria Australia, belonging to two major lithologies, sedimentary limestone and igneous volcanic rock: 3M-3 – Shades of death (S 37.40591; E 148.21114); 3M-49 – Scrubby creek (S 37.44000; E 148.17000); 3H-11 – Harman 1 (S 37.9075; E 141.9769); and 3H-9 – Tunnel cave (S 38.05706; E 141.92000). Two limestone caves (M3 / M49) were sampled in April 2021 and two volcanic caves (H-11 / H-9) were sampled in November 2021. Samples were collected using sterile techniques along a distance transect traversing four sites consisting of the cave entrance at the surface (Entrance) and cave interior (Twilight, Transition and Deep), with the Deep sites representing the furthest point from the entrance (Fig. 1a). At each site (~25 m^2^), three sampling plots (~1 m^2^) were selected at random (Fig. 1a). At each plot, soil-atmosphere fluxes were measured, and samples of soils and biofilms were collected in triplicate for *ex situ* oxidation measurements, physicochemical analysis, and DNA extractions to perform qPCR and metagenomic sequencing. Additionally, opportunistic sampling of conspicuously hued biofilms, fungal spores, moonmilk^1^ and soils were collected for metagenomic sequencing. An alternative design was used for Harman 1 cave, given the lack of bulk soils on the cave floor and high prevalence of brown biofilms meant flux measurements could not be taken. Instead, sampling focused on ambient air measurements and the collection of soils from three plots at the Entrance and biofilm from three plots at the Deep zone and two plots at the Transition site. All sampling occurred during daylight hours and dry weather conditions, gas samples and incubations were processed within 48 h of collection, and soil samples for DNA extraction were initially stored at 5 °C for transport and frozen at -20 °C until DNA extraction.

#### **Soil physicochemistry analysis**

For soil chemistry analysis, surface soils samples from each plot were pooled to form one representative composite sample per site and sent to the Environmental Analysis Laboratory (EAL), Southern Cross University. In total, 37 separate soil physicochemical parameters were selected for analysis, based on commonly reported drivers of soil microbial composition globally. These included: Phosphorus (mg/kg P), Nitrate Nitrogen (mg/kg N), Ammonium Nitrogen (mg/kg N), Sulfur (mg/kg S), pH , Electrical Conductivity (dS/m), Estimated Organic Matter (% OM), Exchangeable Calcium (mg/kg), Exchangeable Magnesium (mg/kg), Exchangeable Potassium (mg/kg), Exchangeable Sodium (mg/kg), Exchangeable Aluminium (mg/kg), Exchangeable Hydrogen (mg/kg), Effective Cation Exchange Capacity (ECEC) (cmol+/kg) , Calcium (%) , Magnesium (%), Potassium (%), Sodium - ESP (%), Aluminium (%), Hydrogen (%), Calcium/Magnesium Ratio, Zinc (mg/kg), Manganese (mg/kg), Iron (mg/kg), Copper (mg/kg), Boron (mg/kg), Silicon (mg/kg Si), Total Carbon (%), Total Nitrogen (%), Carbon/Nitrogen Ratio, Chloride Estimate (equiv. mg/kg), Total Organic Carbon (%), Moisture content (%), Gravel (%), Sand (%), Silt (%), Clay (%).

#### ***In situ* soil–atmosphere gas fluxes**

*In situ* soil-atmosphere fluxes of H_2_, CO, and CH_4_ were measured using static flux chambers. The chamber consisted of a polyvinylchloride (PVC) pipe of 20 cm height and 15 cm diameter, with a threaded access cap. The cap was fitted with a gastight O-ring, two butyl rubber septa (one for air sampling and one for a thermometer), and an axial fan on the inside to promote internal mixing. At each plot, a PVC base collar of 10 cm height and 14.8 cm diameter was inserted ~5 cm into the soil and left to equilibrate for ~15 minutes prior to sampling to reduce lateral gas fluxes. Once the chamber was fitted over the collar, the cap was closed and the axial fan was started. The first measurement taken within ~30 seconds and six measurements were taken at five-minute intervals. Additionally, ambient air measurements were taken over a 30-minute period. For each measurement, 25 mL of gas was collected using a gas tight Terumo syringe fitted with a Luer Lock and Discofix three-way stopcock and measured by gas chromatography as described below. Control gas measurements of ambient air were taken directly before, during, and after sampling. The temperature of the chamber, ambient air, and soil were monitored throughout. Concentrations were then converted to nmol m^-3^ at ambient pressure and temperature using the ideal gas law. Atmospheric flux (*J*_atm_) was calculated from the initial ambient air concentration and rate change at chamber deployment, using linear regression and an exponential model for each chamber measurement as previously described^2^.

#### ***Ex situ* bulk soil oxidation rates**

Soil microcosms were used to determine the capacity of soil microbial communities to oxidize H_2_, CO, and CH_4_ by gas chromatography. Overall, 44 biological replicates from four caves consisting of the Entrance (n = 12), Twilight (n = 9 plots), Transition (n = 11) and Deep (n =12) zones were used. 5 g of soil / brown biofilm (Harman 1) samples were placed in a 120 ml serum vial and incubated at 20°C. The ambient air headspace was amended with H_2_, CO, and CH_4_ (via a mixed gas cylinder containing 0.1 % v/v H_2_, CO, and CH_4_ each in N_2_, BOC Australia) to give starting mixing ratios of approximately 10 parts per million (ppmv) for each gas. At each time interval, 2 ml of headspace gas was sampled using a gas-tight syringe and stored in a sealed 3 ml glass exetainer flushed prior with ultra-high purity N_2_ (99.999% pure, BOC Australia). A VICI gas chromatograph machine with a pulsed discharge helium ionization detector (model TGA-6791-W-4U-2, Valco Instruments Company Inc.) and an autosampler was used to measure gas concentrations as previously described^3^. The machine was calibrated against ultra-pure H_2_, CO and CH_4_ standards down to the limit of quantification (H_2_: 20 ppbv; CO: 9 ppbv; CH_4_: 500 ppbv). Calibration mixed gas (10.20 ppmv of H_2_, 10.10 ppmv of CH_4_, 9.95 ppmv of CO in N_2_, Air Liquide Australia) and pressurized air (Air Liquide Australia) with known trace gas concentrations were used as internal reference standards. Pooled heat-killed soils for each triplicate (treated at 121°C, 15 p.s.i. for 60 mins) and empty vials (duplicate) were prepared as negative controls. First order reaction rate constants were calculated by fitting an exponential model as determined by the lowest overall Akaike information criterion value when compared to a linear model.

#### **Cell specific power calculations**

The calculation was performed as previously described^4,5^, involving the determination of cell-specific power generated by the oxidation of H_2_, CO, and CH_4_, measured as Gibbs energy per unit time per microbial cell. This was accomplished by factoring in the reaction rate for each gas, Gibbs energy of the reaction, and the number of microbial cells involved. The Gibbs energy calculations considered the standard Gibbs energy, the reaction quotient, gas-phase compound activities, and soil conditions. Measurement of reactant concentrations and reaction rates was conducted via gas chromatography. Estimates of microbial cells performing these reactions, were calculated from 16S rRNA gene copy number datasets, and abundances inferred from metagenomics datasets, focusing on trace gas oxidising communities. This approach was compared to previous literature by recalculating the theoretical maximum population of trace-gas oxidizing bacteria using modelled Gibbs energy and measured reaction rates.

#### **Ammonium and sulfide oxidation**

Oxic slurry experiments were undertaken to determine the oxidation rates of ammonium and sulfide of the cave sediments. Biotic and abiotic oxidation rates were distinguished using sterilised (autoclaved at 120°C for 1 hour) and non-sterilised cave sediments. Slurries containing 10 g sediment (wet weight) and 200 mL substrate amended-ultrapure water (50 uM of either ammonium chloride or sodium sulphide) were prepared in 250 mL Schott bottles. The slurries were aerated for 5 minutes to ensure oxic conditions. The bottles containing the slurries were left uncapped but loosely covered with pre-combusted aluminium foil. The slurries were incubated in the dark and were mixed from time to time for the duration of the incubation period (up to 22 days). At each time point, 15 mL of samples were collected and filtered through 0.22 µm pore-sized filters (Sartorius Minisart syringe filter). Filtered samples were analysed for ammonium, nitrate, nitrite and sulfate concentrations using a Lachat Quickchem 8000 Flow Injection Analyzer (APHA standard method). Oxidation rates of ammonium were calculated using linear and non-linear regression of uptake over time as well as liner regression of nitrite and nitrate accumulation over time. Similarly, rates of sulfide oxidation were calculated from linear regression of sulfate increase over time.

**Measurement of carbon isotope at natural abundance level (δ13C)**

Stable isotope measurements were performed to probe possible pathways contributing to the formation of organic matter in caves. Approximately 1 g of sediment was dried at 60°C for 24 h, pulverised and	homogenis	ed with a clean mortar and pestle. The pulverised	sample was further dried at 60°C overnight. Biofilm samples were first resuspended in sterile ultrapure	water, pelleted by centrifugation at 13500 *g* for 2 min, and dried at 60°C. Samples were weighed into	8 mm × 5 mm silver capsules	, repeatedly treated with 10% hydrochloric acid (HCl) to remove inorganic carbon, and dried at 60°C until no further effervescence was observed. All samples were analysed on an ANCA GSL2 elemental analyser	interfaced	to a Hydra 20-22 continuous flow isotope ratio mass spectrometer (CF-IRMS; Sercon Ltd. UK). The precision of the analysis was ±0.2‰. To ensure the accuracy of the isotopic values, internal standards (i.e. sucrose, gelatine and bream) were run concurrently with the samples. These internal standards have been calibrated against internationally recognised reference materials (i.e. USGS 40, USGS41, and IAEA C-6). In addition to δ13C, t	he instrumental analysis also yielded information o	n the percentage of organic carbon	in each	sample, with a	quantification limit of 0.1 mg and	a precision of 0.5 µg for carbon content	.

**Culture-based natural isotope preparation**

To contextualize the δ13C of biomass originating from different carbon fixation pathways, we referred to the carbon isotopic fractionation effects observed in pure cultures. Our dataset includes previously reported literature values for autotrophic pure cultures^6–10^ and new analyses of two hydrogenotrophs (*Cupriavidus necator* H16, *Hydrogenophaga pseudoflava* 1034) and three methanotrophs (*Methylocapsa gorgona* MG08, *Methylococcus capsulatus* NCIB 11132, *Methylosinus trichosporium* OB3b). *H. pseudoflava* and *C. necator* grew autotrophically (CBB cycle) on H_2_ as follows. Pre-culture of both species was prepared by cultivating autotrophically on 20% H_2_ + 5% CO_2_ in minimal medium DSMZ 81 with an initial OD_600_ of 0.03 in gas tight 120 ml serum vials sealed with rubber stoppers and aluminium caps at 30°C with an agitation speed of 180 rpm. The main cultures were prepared by inoculating with an initial OD_600_ of 0.05 under identical conditions in a 1L Schott bottle, sealed with rubber stoppers (GL 45 open-top cap) and screw caps (GL 45 cap open-top). *M. gorgona*, *M. capsulatus*, *M. trichosporium* were grown on CH_4_ as follows. NCIMB nitrate mineral salts (NMS) medium 131 was used for *M. capsulatus* and *M. trichosporium* while DSMZ medium 921 supplemented with 10 ml/L of 10x NMS salts and lanthanum (a final concentration of 1 µM) was used for *M. gorgona*. Exponential *M. capsulatus* and *M. trichosporium* cultures were grown in 1L Schott bottles under 20% v/v CH_4_ in air at 37°C and 30°C, respectively, with shaking (200 rpm) for 7 days. *M. gorgona* cultures were prepared by inoculating 250 mL medium with 70 mL of an early stationary phase culture in 1L glass bottles under 30% v/v CH_4_ in air at 20°C without shaking. δ^13^C values of CO_2_ and CH_4_ were determined to be -10.6 ± 0.1‰ and -51.9 ± 1.5‰, respectively. Triplicate cultures were prepared for all strains. Following incubation, cells at late exponential or stationary phase were subject to centrifugation at 4°C. Supernatants were discarded and cell pellets were resuspended in 1x PBS buffer, followed by another round of centrifugation. Supernatants were discarded again and the cell pellets were stored at -80°C until freeze-drying. Cell pellets were lyophilized for 48 hr and analysed for δ13C, using the same method as for the sediment and biofilm samples. The expected δ^13^C values of autotrophs reported in literatures and current measurements when grown on atmospheric CO_2_ (δ^13^C average -8.5‰)^11^ and CH_4_ (δ^13^C average -47.2‰)^12^ were reported. A random forest model was then generated assessing the importance of key functional genes (pmoA, mcr, hbsT, aclB and rbcL) against depletion of biomass ^13^C stable isotope (δ^13^C_organic_), using the *randomForest* function. Number of variables randomly sampled as candidates at each split = 10, number of trees grown = 10000 and sampling of cases was conducted with replacement.

#### **^14^C isotope labelling**

A radiolabelled carbon dioxide (^14^CO_2_) incubation assay was used to measure the capacity of cave soils / biofilms to mediate three processes: (i) dark CO_2_ assimilation / fixation, (ii) hydrogenotrophic CO_2_ fixation, and (iii) photosynthetic CO_2_ fixation as previously reposrted^13–16^. The sample was added with 50% v/w of sterile water containing 0.0833 μmol of NaH^14^CO_3_ (equivalent amount as 400 ppmv headspace ^14^CO_2_; Perkin Elmer, 56.6 mCi mmol^-1^) and the vial was immediately sealed with PTFE/silicone septum lid (Supelco, Sigma-Aldrich). The samples were then incubated for 96 h at either 20 °C under three conditions: 1) dark (covered in aluminium foil), 2) light (40 μmol photons m^-2^ s^-1^ under constant illumination), and 3) dark hydrogenotrophic condition (100 ppmv of headspace H_2_). For each condition and location, a technical triplicate and a heat-killed soil control was prepared. Following incubation, each sample was treated with 2 ml of 1M HCl to remove unfixed CO_2_ and the content was transferred to a 20-ml scintillation vial. After overnight acidification treatment, the sample was added with an additional 1 ml of 1M HCl and allowed to dry at 60°C under a heat lamp. 20 ml of liquid scintillation cocktail (EcoLume™, MP Biomedical) was added to the dried sample and the signal of fixed ^14^C was measured on an automated liquid scintillation counter (Tri-Carb 2810 TR, Perkin Elmer) for 5 min. The machine was regularly calibrated with ^14^C standards of known activity.

#### **Community DNA extraction**

Total community DNA was extracted from a total of 105 samples using 0.25 g of sediments and biofilms. Extractions were performed using the FastDNA Spin Kit for Soil according to the manufacturer’s instructions with an additional round of bead beating to improve DNA yields. Samples were eluted in DNase- and RNase-free UltraPure Water (ThermoFisher). A sample-free negative control was also extracted. Nucleic acid purity and yield were measured using a NanoDrop ND-1000 spectrophotometer and a Qubit Fluorometer 2.0.

**Quantitative PCR**

Quantitative polymerase chain reactions (qPCR) were used to estimate total bacterial and archaeal biomass of biocrust and topsoil samples. The 16S rRNA gene was amplified using the degenerate primer pair (515F 5ʹ-GTGYCAGCMGCCGCGGTAA-3ʹand 806R 5ʹ-GGACTACNVGGGTWTCTAAT-3ʹ). A synthetic *E. coli* 16S rRNA gene sequence in a pUC-like cloning vector (pMA plasmid; GeneArt, ThermoFisher Scientific) was used as a standard. PCR reactions were set up in each well of a 96-well plate using LightCycler 480 SYBR Green I Master Mix. Each sample was run in triplicate and standards in duplicate on a QuantStudio 7 Flex Instrument (Applied Biosystems). The qPCR conditions were as follows: pre-incubation at 95 °C for 3 min and 45 cycles of denaturation 95 °C for 30 s, annealing at 54 °C for 30 s, and extension at 72 °C for 24 s. 16S rRNA gene copy numbers were calculated based on a standard curve constructed by plotting average Ct values of a serial dilution of the plasmid-borne standard against their copy numbers.

#### **Metagenome sequencing, assembly, and binning**

Metagenomic shotgun libraries were prepared using the Nextera XT DNA Sample Preparation Kit (Illumina Inc., San Diego, CA, USA) and subject to paired-end sequencing (2 × 150 bp) on an Illumina NovaSeq6000 platform at the Australian Centre for Ecogenomics (ACE), University of Queensland. Raw metagenomic sequences were subjected to quality filtering using the BBDuk function of the BBTools v38.80 (https://sourceforge.net/projects/bbmap/); contaminating adapters (k-mer size of 23 and hamming distance of 1), PhiX sequences (k-mer size of 31 and hamming distance of 1), and bases from 3’ ends with a Phred score below 20 were trimmed. After removing resultant reads with lengths shorter than 50 bp, 93% high-quality read pairs were retained for downstream analysis. Coassemblies were conducted according to cave type (limestone vs basalt) and sample nature (dividing by biofilm, sediment, and moonmilk) were conducted using quality-controlled reads in megahit (v1.2.9) using kmers of 27, 37, 47, 57, 67, 77, 87, 97, 107, 117, and 127. In addition, assemblies for each individual sample were performed using metaSPAdes (v3.15.5) with the same set of kmers^17^. Contigs from both coassemblies and individual assemblies with minimum length of 500 bp were used for binning with MetaBAT2 (v2.12.1)^18^, MaxBin2 (v2.2.7)^19^ and CONCOCT (v1.1.0)^20^ to reconstruct metagenome-assembled genomes (MAGs), which is implemented in the ‘binning’ module of MetaWRAP (v1.3.2)^21^. Subsequently, the ‘bin refinement’ modules of MetaWRAP and DAS tool^22^ were used in parallel to consolidate the MAGs from the three binners into one bin set for each assembly. A final MAG set was obtained by dereplicating MAGs from all assemblies at 99% average nucleotide identity using dRep ‘dereplicate’ (v3.4.2)^1323^. The completeness and contamination of the MAGs were estimated using CheckM2 (v0.1.3)^24^ ‘predict’. Taxonomy of each MAG was assigned using GTDB-Tk ‘classify_wf’ (v2.3.0)^25^ against GTDB release 214. CoverM (v0.6.1) ‘genome’ was used to calculate the relative abundance of each MAGs within each sample

#### **Bacterial and archaeal community profiling**

To profile bacterial, archaeal, and eukaryotic community composition based on the metagenomes, quality-filtered short reads encoding 16S rRNA genes were retrieved and assigned using PhyloFlash v.3.0^26^. The function *PhyloFlash.pl* was given a -taxlevel 6 flag for bacteria/archaea specifying a taxonomic assignment down to genus level. Retrieved sequences were then clustered using *PhyloFlash.pl* into NTUs, mapped to the SILVA database (release 138)^27^, and filtered to exclude rare hits (min. read count ≥ 10). Alpha and beta diversity were calculated using R (version 4.1.0 (2021-05-18) -- "Camp Pontanezen") and the Phyloseq package^28^. First, reads were normalized using the coverage-based rarefaction and extrapolation method implemented in the R package iNEXT^29^. Coverage was calculated for each sample using the function *phyloseq_coverage,* followed by rarefaction of all reads, using the function *phyloseq_coverage_raref* with default parameters. Observed and estimated richness was calculated using the *estimate_richness* function specifying with “observed” and “chao1” flags. Generalised linear models (GLM) were used to test for significant differences in richness between and within caves. Following mean–variance comparisons to detect overdispersion and comparing optimal model fits against the residual plots of each distribution (Poisson, quasipoisson, negative binomical), GLMs were fitted with a quasipoisson distribution. To calculate beta diversity, sample counts were first transformed to relative abundance and clustered using a Non-metric Multidimensional Scaling ordination *via* the *ordinate* function with flags “NMDS” and “Bray” function. A stress plot was used to determine linear (R^2^=0.839) and non-metric (R^2^=0.967) fit between ordination distance and dissimilarity (Bray-Curtis). The final stress of <0.2 (0.182) indicated a good representation in reduced dimension. To test for significant differences in microbial community structure between sites, caves, and lithology a Permutational Multivariate Analysis of Variance Using Distance Matrices (PERMANOVA) was used via the *adonis* function, including and assessment of dispersion via the *betadisper* function using the R package Vegan^30^.

#### **Metagenomic functional analysis and metabolic reconstruction**

To estimate the metabolic capability of the soil communities, quality filtered and unassembled short reads were searched against custom protein databases of representative metabolic markers using DIAMOND v.0.9.31 (query cover > 80%)^31^. Searches were carried out using all quality-filtered unassembled reads with lengths over 120 bp. The metabolic markers searched are involved in oxidative phosphorylation (AtpA), NADH oxidation (NuoF), aerobic respiration (CoxA, CcoN, CyoA, CydA), formate oxidation (FdhA), arsenic cycling (ARO, ArsC), selenium cycling (YgfK), sulfur cycling (AsrA, FCC, Sqr, DsrA, Sor, SoxB), nitrogen cycling (AmoA, HzsA, NifH, NarG, NapA, NirS, NirK, NrfA, NosZ, NxrA, NorB), iron cycling (Cyc2, MtrB, OmcB), reductive dehalogenation (RdhA), phototrophy (PsaA, PsbA, energy-converting microbial rhodopsin (RHO)), methane cycling (McrA, MmoA/MmoX, PmoA), hydrogen cycling (large subunit of NiFe-, FeFe-, and Fe-hydrogenases), isoprene oxidation (IsoA), carbon monoxide oxidation (CoxL, CooS), succinate oxidation (SdhA), fumarate reduction (FrdA), and carbon fixation (RbcL, AcsB, AclB, Mcr, HbsT, HbsC)^32–34^. Results were filtered based on an identity threshold of 50%, except for RHO (40%), NuoF, group 4 NiFe-hydrogenases, MmoA, FeFe-hydrogenases, CoxL, AmoA, NxrA, RbcL (all 60%), PsaA (80%), PsbA, IsoA, AtpA, ARO, YgfK (70%), and HbsT (75%). Read counts for each gene were normalized to reads per kilobase per million (RPKM) and average gene copy per organism as previously described ^4,35,36^. Proteins from each MAGs were predicted intrinsically in CheckM^24^. Best gene hits were filtered to retain only those either at least 40 amino acids in length or with at least 80% query or 80% subject coverage^3^. For predicted proteins, the same thresholds were used as above except for AtpA (60%), PsbA (60%), RdhA (45%), Cyc2 (35%) and RHO (30%)^37^.

Metabolic reconstruction for the three most dominant hydrogenotrophs and methanotrophs was performed via annotation of relevant MAGs with DRAM^38^, based on the KEGG database. Pathways were identified through the KEGG pathway database and literature^39^. No MAGs encoded GSH-linked pathway for formaldehyde oxidation, which is consequently omitted from the metabolism figure.

**Phylogenetic analysis**

The evolutionary relationships of maker genes including methane oxidation (PmoA), hydrogen oxidation (NiFe), CO oxidation (CoxL), ammonia oxidation (AmoA), nitrite oxidation (NxrA), carbon fixation (HbsT, RbcL, AclB), sulfide oxidation (Sqr), thiosulfate oxidation (SoxB) photosynthesis (PsaA, PsbA, RHO) were visualised by constructing protein trees. Briefly, protein sequences retrieved from the MAGs and metagenomic assembled reads by homology-based searches against a subset of reference sequences from a custom database^40^, and were aligned using MAFFT^41^ with default parameters, and subsequently trimmed using Trimal v1.2rev59 (“-gappyout”)^42^. The trimmed multiple sequence alignments were used to infer phylogenetic trees with the maximum likelihood method incorporating substitution model selection and 1000 bootstrap iterations implement in IQ-TREE2 v1.6.12 (“-alrt 1000 -B 1000 -m TEST”) ^43^. Tree rendering was performed using the Python package ETE3^44^ combined with visualisation in ITOL^45^.

#### **Methanotroph phylogenetic and metabolic analysis**

GraftM v 0.14^46^ was used to ascertain gene phylogeny. PmoA and MmoX packages generated by^47^ were used and updated with sequences published in^4,5,48–54^. HMM and DIAMOND (hmmsearch+diamond) search methods were applied, along with a conditional E-value (c-Evalue) threshold of 1e-10^26 46^. Multiple sequence alignment of protein sequences was performed with MAFFT v 7.490^41^, trimming at minimum 20% representation was performed with TrimAl v.1.4.1^42^. Phylogenetic trees of protein sequences were generated with IQ-TREE v. 2.2.0.3^43^ with the ultrafast bootstrap approximation option using 1000 iterations and enabling the ModelFinder option (Best-fit model: LG+F+R6). The trees were rerooted and grouped in ARB v. 6.0.3^55^. Visualisation of protein trees was done in iTOL v6^45^. The genome phylogenetic tree was generated by extracting the multiple sequence alignment of high-quality MAGs (completion >90% and contamination <5%) and known methanotrophs generated by GTDB-Tk (v2.1.1)^25^ against GTDB release 214. The tree was constructed with IQ-TREE v. 2.2.0.3^43^ using model WAG+G and ultrafast bootstrap approximation option using 1000 iterations. The trees were rerooted and grouped in ARB v. 6.0.3^35^ and visualised in iTOL v6^45^. Annotation was performed using DRAM and the KEGG database, and pathways were identified through the KEGG pathway database and literature^39^. No MAGs encoded GSH-linked pathway for formaldehyde oxidation, which is consequently omitted from the metabolism figure.

#### **Methane carbon conversion efficiency**

Theoretical estimation of carbon assimilation into microbial biomass was based on measured methane oxidation rates. Three seminal papers on methane carbon assimilation were used to calculate a constant conversion factor^56–58^. The percentage of methane carbon assimilation varied between 20% and 62%. By utilizing the median for each range and computing the mean (40%), we determined the amount of methane carbon assimilated in units of nmol C cell^-1^ min^-1^ by multiplying the cell specific methane oxidation rate by a constant fraction of 0.4. This measurement provides a quantifiable indicator of the role of aerobic methanotrophs in the carbon cycle. One drawback of this approach is that it does not consider variations in assimilation across different cells and variations in assimilation across different physicochemical conditions.

### **Extended Data Figures**

**Extended Data Figure 1**. Maximum-likelihood phylogenetic tree of carbon monoxide dehydrogenase large subuit (CoxL) amino acid sequences, rooted at midpoint. The tree shows sequences from this study’s MAGs alongside (red) representative reference sequences (black). The tree was inferred using LG+F+I model post selection, used all sites, and was bootstrapped with 1000 replicates.

**Extended Data Figure 2.** Maximum-likelihood phylogenetic tree of [NiFe] hydrogenase amino acid sequences, rooted at midpoint. The tree shows sequences from this study’s MAGs alongside (red) representative reference sequences (black). The tree was inferred using Q.pfam+I+ G4 model post selection, used all sites, and was bootstrapped with 1000 replicates.

**Extended Data Figure 3.** Methanotroph phylogeny and metabolic potential. **a**, Phylogenetic tree of high-quality MAGs (completion >90% and contamination <5%) encoding metabolic potential for aerobic methane oxidation and known methanotrophs. **b**, protein phylogeny of Proteobacterial PmoA and PxmA. Tree was generated with the ultrafast bootstrap approximation option using 1000 iterations and enabling the ModelFinder option (Best-fit model: LG+F+R6). **c**, Methanotroph MAG annotation to identify metabolic pathways and potential for methane carbon assimilation using DRAM and KEGG database..

**Extended Data Figure 4**. Maximum-likelihood phylogenetic tree of ammonia monooxygenase subunit A (AmoA) amino acid sequences, rooted at midpoint. The tree shows sequences from this study’s MAGs alongside (red) representative reference sequences (black) from both ammonia oxidising bacteria (AOB) and archaea (AOA). The tree was inferred using Q.yeast+F+G4 model post selection, used all sites, and was bootstrapped with 1000 replicates.

**Extended Data Figure 5**. Maximum-likelihood phylogenetic tree of nitrite oxidoreductase subunit A (NxrA) amino acid sequences, rooted at midpoint. The tree shows sequences from this study’s MAGs alongside (red) representative reference sequences (black). The tree was inferred using the LG+I+G4 model post selection, used all sites, and was bootstrapped with 1000 replicates.

**Extended Data Figure 6**. Maximum-likelihood phylogenetic tree of Ribulose-1,5-bisphosphate carboxylase/oxygenase (Rubisco) large subunit (RbcL) amino acid sequences, rooted using type II RubisCO sequences (not shown). The tree shows sequences from this study’s MAGs alongside (red) representative reference sequences (black). The tree was inferred using the LG+I+G4 model post selection, used all sites, and was bootstrapped with 1000 replicates.

**Extended Data Figure 7.** The average measurements and standard deviation of in situ fluxes are displayed for each gas along the sampling transect. Before closing the flux chamber, the ambient gas concentration is measured at the first timepoint.

**Extended Data Figure 8.** The average measurements and standard deviation of gas chromatography traces for bulk soil incubations and controls are shown, indicating uptake from ~10 ppmv to below atmospheric levels for each gas (dashed line).

**Extended Data Figure 9.** The measurements of oxic slurry incubations are shown for four nitrogen species and sulfur, representing pooled replicates from three caves.

### **Extended Data Tables**

**Extended Data Table 1 | Soil physicochemistry.** The physicochemical parameters, and measurement units for pooled soil samples across each cave and zone are specified.

**Extended Data Table 2 | Biomass estimates.** DNA concentration, elution volume, sample amount, and 16S rRNA gene copy number normalized to both wet weight and dry weight are shown for each sample used in metagenomic sequencing, including extraction controls.

**Extended Data Table 3 a-g | Taxonomic diversity. a**, Nearest Taxonomic Unit (NTU) table showing quality trimmed counts for each sample. **b**, observed and estimated richness (Chao1) for each sample. **c**, Alpha and beta diversity, testing for significant differences in richness and composition. **d-g**, with relative abundance summaries at phylum, class, order, and genus level across all samples.

**Extended Data Table 4 a-c | Metagenomes and MAGs. a**, Sequence statistics and NCBI accessions for metagenomes of this study and those derived from public data. **b**, Metagenomic short read annotation. **c**. Taxonomy, functional annotation, and relative abundance of metagenome assembled genomes (MAGs).

**Extended Data Table 5 a-c | Gas oxidation rates and fluxes. a**, *In situ* trace gas soil to atmosphere (*j_atm_*) flux summaries. **b**, *Ex situ* trace gas oxidation rates and cell specific power calculations. **c**, Legend for Table 5b.

**Extended Data Table 6 | Oxic slurry nutrient assays.** Uptake and accumulation rates of nitrogen and sulfur species for pooled samples across limestone and basalt caves.

**Extended Data Table 7 | Carbon fixation rates. a,** ^14^C radioisotope labelling data for bulk soils normalised to cell specific rates comparing cave and surface sites. **b**, Estimates of methane carbon assimilation rates derived from *ex situ* studies.

### **Supplementary notes**

**Supplementary Note 1. Phylogeny and physiology of cave methanotrophs.**

Phylogeny of the PmoA sequences, encoded by the commonly used *pmoA* marker gene, indicated that several of the MAGs belong to the atmospheric methane oxidising groups USCα (Extended Data Fig. 7b) and USCγ (Extended Data Fig. 7b). For the gammaproteobacterial MAGs, two MAGs, M3-S_metabat2.bin.345 and H1-B1_maxbin2.bin.5_sub, were classified as novel genera within the order CAJXQU01 (Extended Data Fig. 7a). Only one other genome within this order is in GTDB r214 (GCA_913058255.1, *CAJXQU01 sp913058255,* recovered from a marine metagenome), and does not encode *pmoCAB*. Potentially these genomes are the first representatives of a new methanotrophic order in the Gammaproteobacteria. In both MAGs, PmoA sequences grouped with other gammaproteobacterial PmoA sequences (Extended Data Fig. 7b), however not within the well-defined isolate methanotrophs of the Methylococcaceae and Methylomonadaceae families. H1-B1_maxbin2.bin.5_sub also encoded PxmAB of the *pxmABC* operon, a homologue of the *pmoCAB.* While *pxmABC* is found in many methanotrophs, the function is currently unknown^59,60^. Both MAGs encode *pmoCAB* operon, *xoxF*-type methanol oxidation, and formaldehyde oxidation through the tetrahydromethanopterin (H_4_MPT)-pathway (Extended Data Fig. 7c).

MAG H1-B1_metabat2.bin.32 was classified as a novel species within the gammaproteobacterial family and genus UBA1147 (Extended Data Fig. 7b). Three genomes, of which two encode *pmoCAB*, are present within this family in GTDB r214, (*UBA1147 sp002311315* recovered from a hydrothermal vent metagenome, and *UBA1147 sp024958995,* recovered from a marine metagenome). H1-B1_metabat2.bin.32 encoded a PmoA sequence clustering with Methylococcaceae PmoA sequences, along with two copies of PxmA sequences (Extended Data Fig. 7b). A complete *pmoCAB* operon, along with *xoxF*-type methanol oxidation, and formaldehyde oxidation through the H_4_MPT-pathway was encoded (Extended Data Fig. 7c). SD8037_metabat2.bin.6 and SD8020_metabat2.bin.3 were classified as novel species within the gammaproteobacterial genus USCγ-Taylor (Extended Data Fig. 7b), both encoding PmoA sequences clustering with other USCγ sequences (Extended Data Fig. 7b). Both encode full *pmoCAB* operons along with *xoxF*-type methanol oxidation and formaldehyde oxidation through the H_4_MPT-pathway (Extended Data Fig. 7c). Calcium-dependent methanol oxidation was encoded by *mxaFI* in SD8020_metabat2.bin.3, and partially in SD8037_metabat2.bin.6 (*mxaF*). All gammaproteobacterial MAGs lacked complete metabolic potential to assimilate carbon. Canonical gammaproteobacterial methanotrophs usually utilise the RuMP cycle. While all MAGs encoded *pfkA/B* and *FBA*, the key genes hexulosephosphate synthase (*hxlA/fae-hps*) and hexulosephosphate isomerase (*hxlB*) were not encoded. Reasons for absence of detection could be genome incompleteness, or the gene is divergent enough from the KO database representatives to not be correctly assigned. An incomplete serine cycle is present, missing the key gene HPR^61^.

Cytochrome *c* oxidase was encoded in all gammaproteobacterial MAGs, and the cytochrome *bd* oxidase was also encoded in USCγ-Taylor and CAJXQU01 MAGs, both cytochromes are characterised by high oxygen affinity^62^. Dissimilatory nitrate reduction (*narGHJI*) was fully encoded in *USCγ-Taylor* and *UBA1147* MAGs, while partially encoded in *CAJXQU01* (*narGH* and *narGHJ*). Assimilatory sulphate reduction (*cysDCHJI*, lacking *cysN*) and partial dissimilatory sulphate reduction (*dsrCEF)*, though lacking the key genes *dsrAB*, was encoded in UBA1147 Three MAGs, SE1530_metabat2.bin.5_sub, SE1504_maxbin2.bin.0, and SE1504_metabat2.bin.11 were classified into three different species (<95% ANI) within the alphaproteobacterial *Methylocella* genus (Extended Data Fig. 7b). This genus contains the well-studied isolate *Methylocella silvestris* and the recent upland soil cluster alpha atmospheric methane oxidising isolate *M. gorgona*. SE1504_maxbin2.bin.0 encoded a PmoA sequence clustering with USCα PmoA, while SE1504_metabat2.bin.11 encoded a partial USCα-clustering sequence (67 aa, denoted by * in Fig.3b & c) located at the edge of a short contig (2890 bp). SE1530_metabat2.bin.5_sub encoded no USCα-PmoA sequence but contained a sequence clustering with alphaproteobacterial PxmA sequences (Fig. 3b). All MAGs encoded XoxF-type methanol oxidation, and formaldehyde oxidation through the H_4_MPT-pathway. While SE1530_metabat2.bin.5_sub, SE1504_maxbin2.bin.0 had the genomic potential to assimilate formaldehyde through H_4_F and serine cycle, bin.11 is missing two key serine cycle genes, *sga* and *hprA* (Extended Data Fig.7), and does not encode the H_4_F pathway, perhaps due to genome incompleteness. Oxidation of formate to carbon dioxide was also encoded in all MAGs. Similar to the USCα isolate *M. gorgona* the CBB cycle was incomplete and missing the RuBisCo genes in all three MAGs in contrast to the USCα-encoding MAGs of ^47^ and other members of the *Methylocella* genus^63.^ Control genome *M. silvestris* encode the high oxygen affinity *cbb_3_*-type and *bd*-type cytochromes, while the *Methylocella* MAGs obtained here encode low oxygen affinity type-*o* cytochromes ^62^.

CAJXQU01 MAGs were amongst the three most dominant predicted methanotrophic taxa in the cave system, along with JACCXJ01 and *Methylocella* (Extended Data Fig. 7a). Carbon and energy acquisition pathways from each MAG in these taxonomic groups were compiled, revealing broadly similar methylotrophic lifestyles (Extended Data Fig. 7c). However, in contrast to CAJXQU01 and JACCXJ01, *Methylocella* MAGs from the caves encoded the complete tetrahydrofolate pathway and serine cycle to assimilate carbon from formate (Extended Data Fig. 7c). Additionally, all three taxonomic groups encoded a complete TCA cycle and glyoxylate shunt, the core 3C glycolysis steps of the Embden-Meyerhof-Parnas pathway, and the pentose phosphate pathway, whilst only *Methylocella* MAGs additionally encoded the Entner-Doudoroff pathway.

In addition to the alpha and gammaproteobacterial likely new methanotrophs, copper membrane bound monooxygenases (CuMMOs), representing methane monooxygenase homologs, were identified in the MAGs SD8061_maxbin2.bin.11_sub and M3-B2_maxbin2.bin.72_sub and were classified as novel genera within the Bin18 family (Extended Data Fig. 7c). Both MAGs contained PmoA-like sequences grouping with other PmoA-like sequences from the Binatia class (Extended Data Fig. 7b). M3-B2_maxbin2.bin.72_sub also contained an additional MMO-like sequence that grouped with other hydrocarbon monooxygenases. None of the members within Binatia have been experimentally confirmed to be able to carry out methanotrophy. Furthermore, genes for methanol and formaldehyde oxidation could not be identified, indicating that the Bin18 MAGs are most likely not capable of methanotrophy and may be oxidising other hydrocarbons.

**Supplementary Note 2. Etymological information.**

**‘*Candidatus* Hydrogenomurus’**

We propose renaming the Pseudonocardiaceae genus GCA-003244245 as ‘*Candidatus* Hydrogenomurus’, based on the high-quality annotated genome SE1519_metabat2.bin.2 (completeness/contamination: 100%/2.79%) from Tunnel basalt caves (herein ‘*Candidatus* Hydrogenomurus basanatii’)

Species: ‘*Candidatus* Hydrogenomurus basanatii’ (ba.sa.na’tii. N.L. gen. n. *basanatii*, of the basalt cave (Tunnel Cave), where the bacterium was identified)

Genus: ‘*Candidatus* Hydrogenomurus’ (Hy.dro.ge.no.mu.rus. N.L. neut. n. *hydrogenum*, hydrogen (that which produces water, so called because it forms water when exposed to oxygen); from Gr. neut. n. *hydôr*, water; from Gr. ind. v. *gennaô*, to produce; L. masc. n. *murus*, wall; N.L. masc. n. *Hydrogenomurus*, a hydrogen-using bacterium from cave wall)

Full Classification:

- Domain: Bacteria
- Phylum: Actinomycetota
- Class: Actinomycetia
- Order: Mycobacteriales
- Family: Pseudonocardiaceae
- Genus: GCA-003244245 (proposed as ‘*Candidatus* Hydrogenomurus’)
- Species: ‘*Candidatus* Hydrogenomurus basanatii’

***‘Candidatus* Hydrogenocavus’**

We propose renaming the Egibacteraceae genus JACCXR01 as ‘*Candidatus* Hydrogenocavus’, based on the high-quality annotated genome M3-B2_metabat2.bin.58 (completeness/contamination: 95.56%/3.3%) from Shades of death limestone caves (herein ‘*Candidatus* Hydrogenocavus calcicola’)

Species: ‘*Candidatus* Hydrogenocavus calcicola’ (cal.ci’co.la. L. masc./fem. n. *calx* (gen. calcis), chalk; L. masc./fem. n. suff. -*cola*, an inhabitant; from L. masc./fem. n. *incola*, dweller; N.L. masc./fem. n. *calcicola*, of the limestone cave (Shades of death), where the bacterium was identified)

Genus: ‘*Candidatus* Hydrogenocavus’ (Hy.dro.ge.no.ca.vus. N.L. neut. n. *hydrogenum*, hydrogen (that which produces water, so called because it forms water when exposed to oxygen); from Gr. neut. n. *hydôr*, water; from Gr. ind. v. *gennaô*, to produce; L. n. cava, cave; N.L. masc. n. *Hydrogenocavus*, a hydrogen-using bacterium from caves)

Full Classification:

- Domain: Bacteria
- Phylum: Actinomycetota
- Class: Actinomycetia
- Order: Euzebyales
- Family: Egibacteraceae
- Genus: JACCXR01 (proposed as ‘*Candidatus* Hydrogenocavus’)
- Species: ‘*Candidatus* Hydrogenocavus calcicola’

**‘*Candidatus* Hydrogenolapales’**

We propose renaming the Actinomycetota order JACCUZ01 as ‘*Candidatus* Hydrogenolapales’, based on the annotated genome SD8021_metabat2.bin.4 (completeness/contamination: 84.11%/1.1%) from the limestone cave Scrubby Creek (herein ‘*Candidatus* Hydrogenolapis speleophilus’)

Species: ‘*Candidatus* Hydrogenolapis speleophilus’ (spe.le.o’phi.lus. N.L. masc. adj. speleophilus, cave-loving, referring to the cave environment where the MAG was identified)

Genus: ‘*Candidatus* Hydrogenolapis’ (Hy.dro.ge.no.la’pis. N.L. neut. n. hydrogenum, hydrogen; L. n. lapis, stone; N.L. neut. n. Hydrogenolapis, a hydrogen-oxidizing bacterium)

Family: ‘*Candidatus* Hydrogenolapaceae’ (Hy.dro.ge.no.la.pa’ce’ae. N.L. neut. n. Hydrogenolapis, a (Candidatus) bacterial genus; suff. -aceae ending to denote a family; N.L. fem. pl. n. Hydrogenolapaceae, family of the genus Hydrogenolapis)

Order: ‘*Candidatus* Hydrogenolapales’ (Hy.dro.ge.no.la’pa.les. N.L. neut. n. Hydrogenolapis, a (Candidatus) bacterial genus; suff. -ales ending to denote an order; N.L. fem. pl. n. Hydrogenolapales, order of the family Hydrogenolapaceae)

Full Classification:

Domain: Bacteria

Phylum: Actinomycetota

Class: Actinomycetia

Order: JACCUZ01 (proposed as ‘*Candidatus* Hydrogenolapales’)

Family: JACCUZ01 (proposed as ‘*Candidatus* Hydrogenolapaceae’)

Genus: JACCUZ01 (proposed as ‘*Candidatus* Hydrogenolapis’)

Species: ‘*Candidatus* Hydrogenolapis speleophilus

***‘Candidatus* Methyloligotrophales’**

We propose renaming the Gammaproteobacterial order JACCXJ01 as ‘*Candidatus* Methyloligotrophales’, based on the high-quality annotated genome SD8037_metabat2.bin.6 (completeness/contamination: 96.44%/0.56%) from Shades of Death limestone caves (herein ‘*Candidatus* Methyloligotropha calcicola’)

Species: ‘*Candidatus* Methyloligotropha calcicola’ (cal.ci’co.la. L. masc./fem. n. *calx* (gen. calcis), chalk; L. masc./fem. n. suff. -*cola*, an inhabitant; from L. masc./fem. n. *incola*, dweller; N.L. masc./fem. n. *calcicola*, of the limestone cave (Shades of death), where the bacterium was identified)

Genus: ‘*Candidatus* Methyloligotropha’ (Me.thy.lo.li.go.tro’pha. N.L. neut. n. *methylum*, from French *méthyle*, back-formation from *méthylène*, coined from Gr. n. *methu*, wine, and Gr. n. *hulê*, wood, referring to the methyl group; N.L. pref. methylo-, pertaining to the methyl radical; Gr. masc. adj. *oligos*, little, few; Gr. masc./fem. adj. *trophos*, feeder, rearer, that which nourishes; N.L. fem. n. *Methyloligotropha*, an oligotrophic methyl-using bacterium)

Family: ‘*Candidatus* Methyloligotrophaceae’ (Me.thy.lo.li.go.tro’pha.ce’ae. N.L. neut. n. *Methyloligotropha* a (Candidatus) bacterial genus; suff. -aceae ending to denote a family; N.L. fem. pl. n. *Methyloligotrophaceae*, family of the genus *Methyloligotropha*)

Order: ‘*Candidatus* Methyloligotrophales’ (Me.thy.lo.li.go.tro’pha’les. N.L. neut. n. *Methyloligotropha* a (Candidatus) bacterial genus; suff. -ales ending to denote an order; N.L. fem. pl. n. *Methyloligotrophales*, order of the family *Methyloligotrophaceae*)

Full Classification:

- Domain: Bacteria
- Phylum: Proteobacteria
- Class: Gammaproteobacteria
- Order: JACCXJ01 (proposed as ‘*Candidatus* Methyloligotrophales’)
- Family: JACCXJ01 (proposed as ‘*Candidatus* Methyloligotrophaceae’)
- Genus: USCg-Taylor (proposed as ‘*Candidatus* Methyloligotropha’)
- Species: ‘*Candidatus* Methyloligotropha calcicola’

**‘*Candidatus* Methylocavales’**

We propose renaming the Gammaproteobacterial order CAJXQU01 as ‘*Candidatus* Methylocavales’, based on the high-quality annotated genome H1-B1_maxbin2.bin.5_sub (completeness/contamination: 97.84%/2.43%) from Harman basalt caves (herein ‘*Candidatus* Methylocavus basanatii’)

Species: ‘*Candidatus* Methylocavus basanatii’ (ba.sa.na’tii. N.L. gen. n. *basanatii*, of the basalt cave (Harman 1), where the bacterium was identified)

Genus: ‘*Candidatus* Methylocavus’ (Me.thy.lo.ca.vus. N.L. neut. n. *methylum*, from French *méthyle*, back-formation from *méthylène*, coined from Gr. n. *methu*, wine, and Gr. n. *hulê*, wood, referring to the methyl group; N.L. pref. methylo-, pertaining to the methyl radical; L. n. cava, cave; N.L. neut. n. *Methylocavus*, a methyl-using bacterium from caves)

Family: ‘*Candidatus* Methylocavaceae’ (Me.thy.lo.ca.va.ce’ae. N.L. neut. n. *Methylocavus* a (Candidatus) bacterial genus; suff. -aceae ending to denote a family; N.L. fem. pl. n. *Methylocavaceae*, family of the genus *Methylocavus*)

Order: ‘*Candidatus* Methylocavales’ (Me.thy.lo.ca.va’les. N.L. neut. n. *Methylocavus* a (Candidatus) bacterial genus; suff. -ales ending to denote an order; N.L. fem. pl. n. *Methylocavales*, order of the family Methylocavaceae)

Full Classification:

- Domain: Bacteria
- Phylum: Proteobacteria
- Class: Gammaproteobacteria
- Order: CAJXQU01 (proposed as ‘*Candidatus* Methylocavales’)
- Family: CAJXQU01 (proposed as ‘*Candidatus* Methylocavaceae’)
- Genus: CAJXQU01 (proposed as ‘*Candidatus* Methylocavus’)
- Species: ‘*Candidatus* Methylocavus basanatii’
